# Action Observation Network activity related to object-directed and socially-directed actions in Adolescents

**DOI:** 10.1101/2020.11.29.402560

**Authors:** Mathieu Lesourd, Alia Afyouni, Franziska Geringswald, Fabien Cignetti, Lisa Raoul, Julien Sein, Bruno Nazarian, Jean-Luc Anton, Marie-Hélène Grosbras

**Affiliations:** Laboratoire de Psychologie (EA 3188), Université de Bourgogne Franche-Comté, Besançon, France; Aix Marseille Univ, CNRS, LNC, Laboratoire de Neurosciences Cognitives, Marseille, France; Univ. Grenoble Alpes, CNRS, TIMC-IMAG, F-38000 Grenoble, France; Aix Marseille Univ, CNRS, Centre IRM-INT@CERIMED (Institut des Neurosciences de la Timone - UMR 7289), Marseille, France

**Keywords:** Action Observation, Adolescence, fMRI, Social actions, Object-direct actions

## Abstract

The Action Observation Network (AON) encompasses brain areas consistently engaged when we observe other’s actions. Although the core nodes of the AON are present from childhood, it is not known to what extent they are sensitive to different action features during development. As social cognitive abilities continue to mature during adolescence, the AON response to socially-oriented actions, but not to object-related actions, may differ in adolescents and adults. To test this hypothesis, we scanned with functional magnetic resonance imaging (fMRI) 28 typically-developing teenagers and 25 adults while they passively watched videos of hand actions varying along two dimensions: sociality (i.e. directed towards another person or not) and transitivity (i.e. involving an object or not). We found that observing actions recruited the same fronto-parietal and occipito-temporal regions in adults and adolescents. The modulation of voxelwise activity by the social or transitive nature of the action was similar in both groups of participants. Multivariate pattern analysis, however, revealed that the accuracy in decoding the social dimension from the brain activity patterns, increased with age in lateral occipital and parietal regions, known to be involved in semantic representations of actions, as well as in posterior superior temporal sulcus, a region commonly associated with perception of high level features necessary for social perception. Change in decoding the transitive dimension was observed only in the latter region. These findings indicate that the representation of others’ actions, and in particular their social dimensions, in the adolescent AON is still not as robust as in adults.

**Significance statement:** The activity of the action observation network in the human brain is modulated according to the purpose of the observed action, in particular the extent to which it involves interaction with an object or another person. How this conceptual representation of actions is implemented during development is largely unknown. Here, using multivoxel pattern analysis of fMRI data, we discovered that, while the action observation network is in place in adolescence, the fine-grain organization of its posterior regions is less robust than in adults to decode the social or transitive dimensions of an action. This finding highlights the late maturation of social processing in the human brain.

## Introduction

When we observe other’s actions a set of brain areas is consistently engaged contributing to our social interactions’ capability. The so-called Action Observation Network (AON) comprises fronto-parietal regions -- traditionally associated with action planning (Gallese et al., 1996; Buccino et al., 2001) – as well as posterior superior temporal sulcus (pSTS) and high-level visual occipito-temporal areas -- traditionally associated with perceptual analyses (Carr, Iacoboni, Dubeau, Mazziotta, & Lenzi, 2003; Downing, 2001; for meta-analyses see Caspers, Zilles, Laird, & Eickhoff, 2010; Grosbras, Beaton, & Eickhoff, 2012) ADD OSTENHORF. The AON supports not only the representation of low-level aspects of an action (e.g., kinematics) but also its high-level aspects (e.g., goal, intention) indexing the abstract or conceptual knowledge about the action observed [WURM; LIGNAU; HAFRI; URGEN]. Notably, a number of empirical and theoretical studies suggest that activity in different subsystems of the AON might be modulated by the social aspect of the observed actions, that is whether they involve another agent or not. For instance, watching point-light displays representing two individuals interacting enhanced the recruitment of the inferior frontal gyrus (IFG), premotor areas, bilateral IPS and the right superior parietal lobe (SPL), as compared to watching the same individuals acting independently (Centelles et al., 2011). Higher activity in fronto-parietal (Oberman et al., 2007; Becchio et al., 2012) and occipitotemporal parts (Saggar et al., 2014; Isik et al., 2017; Wurm et al., 2017; Walbrin et al., 2018; Becchio et al., 2012) of the AON has also been reported when participants observed gestures or object-directed actions performed with a social intent (e.g. communicating or cooperating) as compared to individual-centered actions. Multivoxel patterns and representation similarity analyses have revealed a representation of both the object-directedness (transitivity) and person directedness (sociality) qualification of an action in most part of the AON. Yet results seems to converge to indicate that only in the posterior part these representations, and in particular the social representation, generalize well across a variety of perceptually divergent actions (Wurm and Caramazza, 2019; but see Hafri 2017), and even verbal description (Wurm and Caramazza Nat Communication 19.

The general notion of the existence of a conceptual representation of action content and in particular the social orientation of an action, raises the question of the ontogeny of this representation. This question is even more complex given the late maturation of this part of the brain during adolescence, both in terms of structure (e.g. grey matter density) and functional connectivity patterns, in contrast to premotor regions that seem to mature earlier. HYP If shaped by experience fine grained representation should change during adolescence.

Responses of parts of the AON are present very early in development. Activity during passive observation of other people’s hand actions has been reported in sensorimotor (Shimada and Hiraki, 2006) and temporal areas (Lloyd-Fox et al., 2009) in 5-months old infants using near-infrared spectroscopy. Functional magnetic resonance imaging (fMRI) studies in children (from 7 years old) and adolescents showed that all nodes of the AON are identified during action observation (Ohnishi et al., 2004; Shaw et al., 2011, 2012; Pokorny et al., 2015). Direct comparison with adults showed a lower activity in occipito-temporal areas in children age 7-9 (Morales et al., 2019) and less left-lateralization in occipital regions in a group of 7 to 15 years old (Biagi et al., 2016). So far, however, no study has investigated whether the representation of the *content* of observed actions is the same as in adults.

Yet social perception skills continue to mature especially during adolescence (Scherf et al., 2007; Ross et al., 2014) while social orientation and social cognition also undergo a drastic increase in complexity (Steinberg and Morris, 2001). Besides, structural changes in the AON regions still occur until the end of the teenage years, which suggests also changes in functional organization (Mills et al., 2014). In these regards, we hypothesized that the modulation of the action observation network when the type of action is of social nature might also change. We designed an fMRI paradigm where adolescents (13-17 years old) and adults passively watched short videos of actions that varied in their social or transitive nature. We asked specifically whether, at an age when the overall activity of the AON is adult-like, the local representations of the different conceptual dimensions of action is also already mature. In line with the delayed development of social cognition, we would expect bigger differences within the AON between adolescents and adults for social actions only.

## 2. Material and Methods

### 2.1. Participants

Twenty-eight typically developing adolescents aged from 13 to 17 years (*M*_*age*_ = 15.1, *SD* = 1.26; 13 females; 27 right-handers) were enrolled in the study. They completed the Pubertal Development Scale (PDS; Petersen & Crockett, 1988), a sex-specific eight-item self-report measure of physical development based on Tanner stages (Marshall & Tanner, 1969, 1970). Adolescents answered questions concerning their physical development (e.g. growth in stature, breast development, pubic hair) and on the basis of their answers they were assigned to one of the categories of pubertal status: mid-pubertal (Tanner stage 3, *n* = 9), advanced pubertal (stage 4, *n* = 13), and post-pubertal (stage 5, *n* = 6). Twenty-five adults (*M*_*age*_ = 26.6, *SD* = 2.02, range = 24-33 years old; 14 females; 22 right-handers) were also recruited in the study. Recruitment was made through internal ads in the university.

All participants reported to be healthy and typically developing, they had normal or corrected-to-normal vision and reported no history of neurological or psychiatric disorder. All participants were voluntary and signed written consent. Written consent was also obtained from the adolescents’ parents. The study was in line with the Declaration of Helsinki and was approved by the national Ethics Committee.

Inclusion in the final sample required that head motion during scanning did not exceed 2mm displacement between consecutive volumes on 90% of volumes for each run. One male adolescent was excluded based on this criterion. One adult was also excluded following technical problems during fMRI scanning.

### 2.2. Stimuli

The stimuli consisted of 256 videos, each representing the same scene with two persons, amongst four possible actors, facing each other across a table, seen from the side (i.e. one actor on each side of the screen). Only the arms and hands of the actors were visible. Different objects were placed on the table. Only one of the two actors produced an action with her/his right or left arm. There were no physical contact between the two actors.

We grouped the actions into four classes, based on whether the action depicted involved the other person or not (Social or Non-Social) and whether it involved an object or not (Transitive or Intransitive). We had 64 exemplars of videos for each class that represented the following actions: (1) Social Transitive (ST): give/take pen and give/take book; (2) Non-Social Transitive (NT): write/rub with pencil and open/close book; (3) Social Intransitive (SI): agree/disagree finger gesture and come/go away hand gesture ; and (4) Non-Social Intransitive (NI): stroke/scratch arm with finger and stroke/scratch arm with hand.

To further increase the variability in each class, the action could be performed by the actor sitting on the left or right side of the table and filmed from two slightly different perspectives. This maximized chances to identify representational mechanisms that rely on abstract action representations that generalize across perceptual information (Wurm, Ariani, Greenlee, & Lingnau, 2016; Wurm, Caramazza, & Lingnau, 2017; Wurm & Lingnau, 2015).

In addition, we added control items consisting of eight modified action videos from the four action classes (2 control videos per action class). In these videos, the actors were removed, and a pink disk moved within the scene. The trajectory and cinematic of the disk were matched with that of the gesture from the original video.

All videos had a duration of about 3 seconds (with 30 frames per second) and a resolution of 640 × 480 pixels. All 256 videos were manually inspected with mpv media player (available from https://mpv.io/) to determine the onset and duration of each action. Individual action duration was then standardized across action class by slightly speeding up or slowing down the individual videos for which the duration of the action fell outside the mean +/− two times the standard deviation of all videos of the respective action class. A variable number of ‘filler’ frames before and after the execution of the action were included for each video to create final trial videos, consisting of a combination of three videos of the same action class each (see below), of equal length. All video editing was performed using ffmpeg (version 3.2, available from http://ffmpeg.org/) and in-house Python scripts. The quantity and spatial amplitude of motion was inevitably different for each class of action. For instance, the social action “Thumb down” implies a large gesture of the arm whereas the non-social action “Scratch” implies a local gesture with low arm amplitude. As a consequence, the global and local visual motion was different across classes. In order to quantify and control in subsequent analyses for potentials effects of these interclass differences, we used a program developed in-house in Python with the library OpenCV (Open Source Computer Vision Library; https://opencv.org/) to compute, for each video frame, the number of pixels that changed intensity relative to the preceding frame. Then, the total number of changing pixels was divided by the total number of frames to obtain a score of motion magnitude. Videos of social actions involved more visual motion than videos of non-social actions. We thus used the motion magnitude score as a regressor of non-interest in the analysis of brain activity (see section Univariate fMRI Analysis, for more details).

All videos were tested in a separate online experiment using the platform Testable (https://www.testable.org/). We created 8 subsets of 64 videos where all classes of actions were equally represented. For this experiment, we recruited 126 participants (*M* = 33.9 years, *SD* = 10.2; 77 females) who were randomly assigned to one of the eight subsets of videos and were asked to rate each video using visual analog scales (from 0 = not at all to 100 = very much), along two dimensions introduced with the following questions: for sociality, “How much is the action relevant for the nonacting person?”; for transitivity, “How much does the action involve the interaction with a physical object?”. As expected, the four categories were well-discriminated, even if there was more variability along the social dimension for the transitive actions.

For the fMRI experiment, to maximize the BOLD response elicited by each action observation condition, videos were arranged in triplets that varied across the identity of actors, the perspective, and the side of action. This resulted in 9.5 s videos showing the same action class, hereafter called trial videos, that were used in a block design.

### 2.3. fMRI experiment

Each participant was scanned in a single-session with: (i) a T1-weighted anatomical scan, (ii) one practice functional run to ensure that participants felt comfortable with the task, (iii) eight functional runs. Each functional run contained 20 trials (16 action trials plus 4 control conditions; see **Figure 1**). Each trial started with a fixation cross (variable duration from 1 to 3 s) followed by a trial video (9.5 s), which was then immediately followed by a blank screen (variable duration from 0.5 to 1.5 s) and a subsequent rating screen (5 s). The inter-trial-interval thus varied from 16.12 s to 19.12 s. Each run ended with a 10 s fixation period. A genetic algorithm was used to optimize the experimental design with regards to contrast estimation (Wager and Nichols, 2003; Kao et al., 2009) using the toolbox NeuroDesign (https://neurodesign.readthedocs.io/en/latest/index.html). We thereby created eight different schedules of sequences of conditions and intertrial intervals. The assignment of these schedules to the eight runs was counterbalanced across participants.

**Figure 1.**
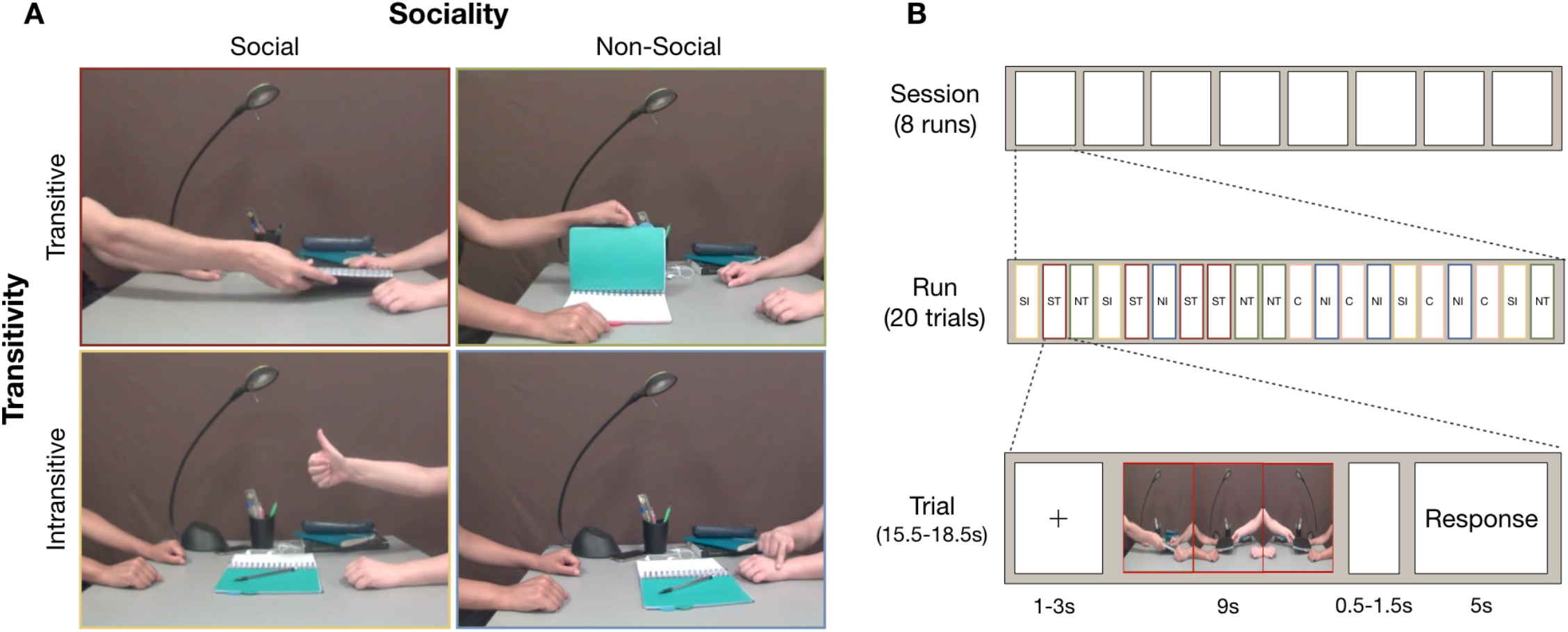
**(A)** stimuli used in the present study varying across two dimensions: sociality (social, non-social) and transitivity (transitive, intransitive), leading to 4 distinct categories of actions: Social Transitive (ST), Non-Social Transitive (NT), Social Intransitive (SI), and Non-Social Intransitive (NI). Each category was made of four classes of actions: (ST): Give: the actor moves a book or a pen from his/her peri-personal space toward the peri-personal space of the passive actor; Take: the reverse of Give; (NT): Open: the actor opens a notebook; Close: the reverse of Open ; Rub: the actor rubs pencil trace on the notebook with rapid oscillatory movements; Write: the actor writes something on the notebook with the pencil; (SI) Agree: the actor indicates with a gesture (i.e., thumb up) to the passive actor that he agrees; Disagree: the actor indicates to the passive actor with a gesture (i.e., thumb down) that he disagrees; Come: the actor indicates with his active hand to the passive actor to come closer; Go away: the reverse of Come; (NI): Stroke: the actor strokes his forearm with his active hand; Scratch: the actor scratches his forearm with his active hand. **(B)** schematic depiction of the sequence of events in a representative session.

In the scanner, stimuli were back-projected onto a screen (60 Hz frame rate, 1024 × 768 pixels screen resolution) via a liquid crystal DLP projector (OC EMP 7900, Epson) and viewed through a mirror mounted on the head coil. Image on the screen had a 40×30 cm size, covering a 20° angle of view. Participants gaze position on the projection mirror was recorded (Eyelink 1000 system, SR Research). Before each functional run, the spatial accuracy of the calibration of the eye tracker was validated using 9 points. If the average deviation exceeded 1° of visual angle, the spatial calibration was redone. Stimulus presentation, response collection and synchronization with the fMRI acquisition triggers and the eyetracker were implemented in a custom-built program, using the LabVIEW (National Instrument) environment. After each functional run, participants were allowed self-determined breaks.

### 2.4. Task

Participants were first asked to watch attentively each trial video. Immediately after a trial video, a response screen, showing a question and a slider, was presented and participants had to indicate, depending on the question, either the degree of sociality or the degree of transitivity of the action that was depicted in all the three videos they had just seen. We used the same questions as in the preliminary independent experiment. Participants gave their response by moving a track-ball with their right index along an analog-scale (from 0 = not at all to 100 = very much) and validated their choice by clicking with their right thumb. Only one question was displayed for each trial. As a trial video was presented twice during the experiment, both social and transitive ratings were collected for each action. The order of presentation of the questions was counterbalanced across subjects. Ratings were used to ensure that adolescents and adults were able to discriminate the items across sociality and transitivity. Importantly, as participants did not know in advance which question would be asked, they were not biased towards attending to one or the other dimension. Two questions were also asked for the control videos, one concerning the distance covered and the other concerning the velocity of the pink disk. To ensure that participants understood and followed correctly the instructions during the fMRI session, they completed a practice run before the scanning, outside the scanner. No information about the exact aim of the study was given before the experiment.

### 2.5. Data acquisition

Imaging data were acquired on a 3T Siemens Prisma Scanner (Siemens, Erlangen, Germany) using a 64-channel head coil. Blood-Oxygen Level Dependent (BOLD) images were recorded with T2*-weighted echo-planar images acquired with the multi-band sequence (version R016a for Syngo VE11B) provided by the University of Minnesota Center for Magnetic Resonance Research (https://www.cmrr.umn.edu/multiband/). Functional images were all collected as oblique-axial scans aligned with the anterior commissure–posterior commissure (AC–PC) line with the following parameters: 287 volumes per run, 54 slices, TR/TE = 1224 ms / 30 ms, flip angle = 66°, field of view = 210 × 210 mm^2^, slice thickness = 2.5 mm, voxel size = 2.5x 2.5 × 2.5 mm^3^, multiband factor = 3. To correct for magnetic field inhomogeneity during data preprocessing, we also acquired a pair of spin-echo images with reversed phase encoding direction (TR/TE = 7.060 ms / 59 ms, flip angle = 90°, voxel size = 2.5 × 2.5 × 2.5 mm^3^). Structural T1-weighted images were collected using a T1 weighted Magnetization-Prepared 2 Rapid Acquisition Gradient Echoes (MP2RAGE) sequence (176 sagittal slices, TR/TE = 5000 / 2.98 ms, TI1/TI2 = 757 / 2500 ms, alpha1/alpha2 = 4° / 5°, Bandwidth = 240Hz/pix, Field-Of-View = 256 × 256 × 176 mm^3^, slice thickness = 1 mm, voxel size = 1 × 1 × 1 mm^3^).

### 2.6. Preprocessing

Structural T1-weighted images were derived from MP2RAGE images by removing the noisy background and were skullstripped and segmented into tissue type (GM: grey matter, WM: white matter and CSF: cerebro-spinal fluid tissues) using the Computational Anatomy Toolbox (CAT12; http://dbm.neuro.uni-jena.de/cat12/). Functional data were analyzed using SPM12 (Wellcome Department of Cognitive Neurology, http://www.fil.ion.ucl.ac.uk/spm) implemented in MATLAB (Mathworks, Sherborn, MA). Preprocessing for univariate analyses included the following steps (1) realignment to the mean EPI image with 6-head motion correction parameters; (2) co-registration of the individual functional and anatomical images; (3) normalization towards MNI template; (4) spatial smoothing of functional images (Gaussian kernel with 5 mm FWHM). For multivariate pattern analyses step (2) and step (3) were skipped to work only on unsmoothed EPI images, in native space of each subject.

### 2.7. Univariate fMRI analysis

A general linear model (GLM) was created using design matrices containing one regressor (explanatory variable) for each condition of interest (i.e., social transitive, social intransitive, non-social transitive, and non-social intransitive) modeled as a boxcar function (with onsets and durations corresponding to the start of each video of that condition) convolved with the canonical hemodynamic response function (HRF) of SPM, one regressor for the control condition, built the same way, one regressor accounting for judgement and motor response (HRF-convolved boxcar function containing all the periods during which the rating screen was presented and responses given) and six regressors of non-interest resulting from 3D head motion estimation (x, y, z translation and three axis of rotation). As quantity and spatial amplitude of visual motion was different for each class of action, we also included one regressor controlling for unequal motion quantity. This regressor was modeled as a boxcar function with onsets and durations of each video convolved with the canonical HRF and parametrically modulated with motion quantity values (*z*-scored for each run). A regressor accounting for eye movements was also included with each saccade modeled according to its onset and duration, convolved with the canonical HRF. In addition, in order to estimate and remove the variance corresponding to physiological noise, we used the PhysIO toolbox (Kasper et al., 2017). We extracted the time-course of the signal from all voxels in the CSF and separately in the white matter. A principal component analysis (PCA) was performed (i.e., CompCor; Behzadi et al., 2007), and fourteen physiological components related to non-BOLD activity were extrapolated in the normalized WM (6 first PCs + mean signal) and in the normalized CSF (6 first PCs + mean signal). We included these fourteen components as confounds regressors in the GLM. The model was estimated in each participant, also taking into account the average signal in each run. The contrast of parameter estimates of each condition compared to control, computed at the individual level, were entered into a three-way repeated measures ANOVA, with Group (Adolescents vs Adults) as between-subject factor, and Sociality (social vs non-social) and Transitivity (transitive vs intransitive) as within-subject factors. The analysis was performed using GLMflex (http://mrtools.mgh.harvard.edu/index.php/GLM_Flex) implemented in Matlab. We present results maps with a significance threshold set at *p*FWE < .05 with family-wise error (FWE) correction applied at the cluster level (cluster-defining non-corrected threshold at *p* < .001).

### 2.8. Multi-voxel pattern analysis

#### 2.8.1. Regions of interest (ROI) definition

In a first analysis, we focused on regions typically recruited during action observation. We defined eight ROIs: bilateral LOTC, bilateral PMv, bilateral pSTS, and bilateral IPS/SPL. These ROIs were derived from an independent meta-analysis of fMRI and PET data (Grosbras et al., 2012), by taking the conjunction of activated voxels reported in a set of studies contrasting observing hand movements (with or without object) to control conditions (*p* < .001 uncorrected, cluster extent threshold of 5 voxels).

All ROIs, defined in MNI-space, were transformed into each subject native space and masked with his grey matter mask. Importantly, overlapping voxels across ROIs were manually inspected and were attributed to the smallest ROI to ensure all ROIs were independent of each other (Bracci et al., 2017). This concerned only left pSTS and left LOTC due to their spatial proximity in the meta-analysis results and represents a marginal number of voxels (*M* = 6.51 ± 2.68).

Each ROI had a different number of voxels across subjects and hemispheres (mean size and standard deviation are indicated in brackets): LOTC (left = 236.45 ± 33.60, right = 316.57 ± 51.57), PMv (left = 139.57 ± 23.15, right = 143.06 ± 24.77), pSTS (left = 71.84 ± 13.20, right = 75.43 ± 17.98), and IPS/SPL (left = 213.04 ± 37.92, right = 171.92 ± 27.93). These differences may prevent the reliability of between subjects’ comparisons and may bias group comparisons. To obtain the same sizes across participants, we applied a voxel selection procedure, separately for each classification analysis, based on the highest values in the univariate F-test. Thereby for each subject we defined ROIs with size constrained by the smallest size observed across participants in the initial definition: LOTC [*n* = 174 voxels], PMv [*n* = 96 voxels], pSTS [*n* = 47 voxels], and IPS/SPL [*n* = 104 voxels].

#### 2.8.2. ROI-based MVPA

We performed multivoxel pattern analyses (MVPA) within the eight ROIs independently. At the individual level we computed a new GLM using the realigned and unwarped images in native space (without smoothing) and estimating single trial activity (i.e. using 20 regressors per run). The new GLM included the same covariates used in the univariate analysis. In total this procedure resulted in 32 maps of parameter estimates (beta) per action condition (4 action exemplars x 8 runs) for each subject (total 128 maps). MVPA was performed using nilearn (Abraham et al., 2014) for Python 3.7. For voxels within each ROI, we trained, on a subset of data, a linear support vector machine classification (regularization hyperparameter C = 1), to distinguish patterns of parameter estimates associated with each condition. We then tested the classifier ability to decode the conditions associated with patterns of parameter estimates on the remaining data. We used an eight-fold leave-one out cross-validation schedule, training on data from seven runs and testing on data from the remaining run and averaging the classification accuracies (percent correct) across the eight iterations.

This procedure was carried out independently in each ROI in two analyses: firstly, we trained the classifier to discriminate social versus non-social actions (112 patterns from the seven runs of the training set: 56 social and 56 non-social), independently of the transitive dimension, and we tested on the remaining 16 patterns (8 social and 8 non-social); secondly, we trained the classifier to discriminate transitive versus intransitive actions (112 patterns: 56 transitive and 56 intransitive), independently of the social dimension, and we tested on the remaining 16 patterns (8 transitive and 8 intransitive) (see **Figure 2**). For each analysis, to make group-level inferences we compared the averaged accuracies per ROI to chance level (50%) using a one-tailed one-sample Student *t*-test. Statistical results were FDR-corrected for the number of ROIs (Benjamini and Yekutieli, 2001). We also assessed the significance of decoding at the individual level with a fold-wise permutation scheme (Etzel and Braver, 2013). To do so, the classification was repeated 1000 times after randomizing the labels in order to construct a null-distribution per subject, ROI and condition. The *p*-value was then given by dividing the number of times where the mean classification accuracy was greater than the classification score obtained by permuting labels, by the number of permutations.

**Figure 2.**
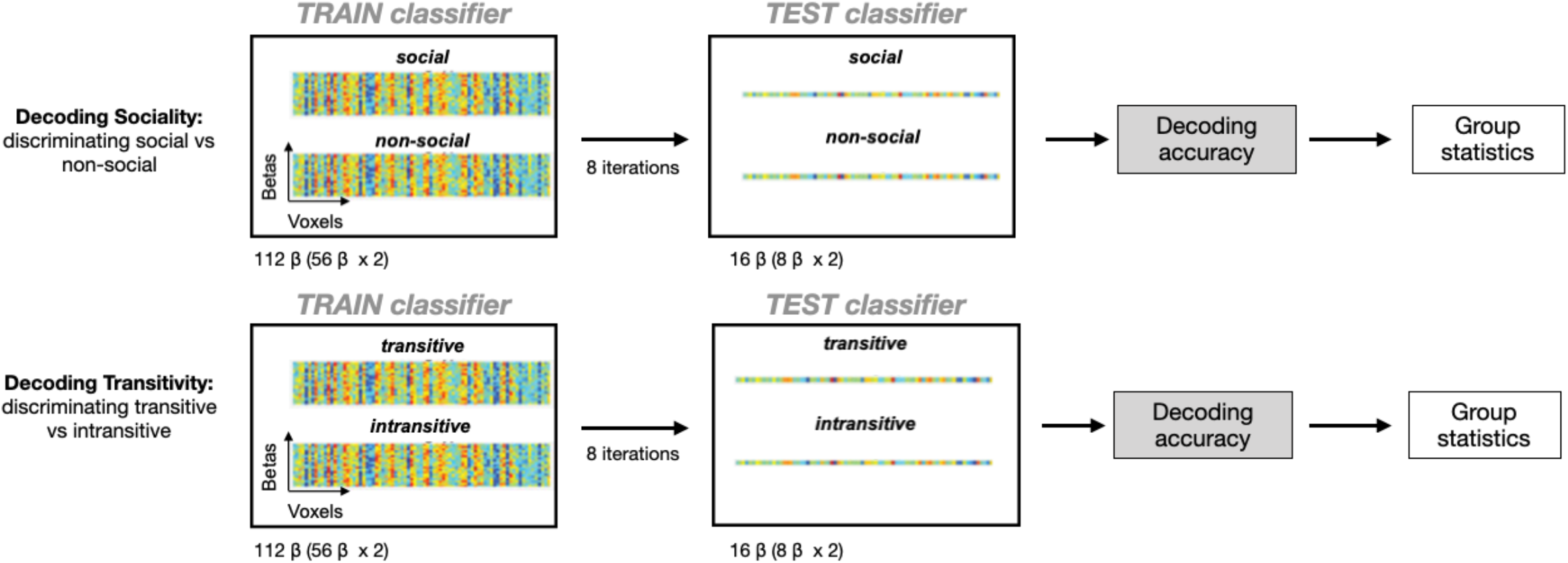
Schematic representation of the MVPA. A beta estimate was first extracted for each trial using a GLM. The SVM classification was performed using a leave-one-out cross-validation scheme. A SVM classifier was trained (112 β) and tested to discriminate between Social versus Non-Social actions (16 β) and between Transitive versus Intransitive (16 β) actions. Classification accuracies were averaged across iterations (8 iterations) and entered in a two-way ANOVA with Age group (Adolescents and Adults) as between factor and hemisphere (Left and Right) as within factor, for each ROI and each Action Dimension separately.

We entered classification accuracies in a two-way ANOVA with Age group (adolescents and adults) as between factor and Hemisphere (Left and Right) as within factor.

Finally, we carried out a four-way classification (NI, NT, SI, ST) and built confusion matrices to explore which conditions might be confounded to each other. Mean accuracies for each action class (values in the confusion matrix diagonal) was then entered in ANOVAs to investigate potential difference between adolescents and adults. Additionally, to probe developmental effect, correlations between mean accuracy score for each action category (ST, SI, NT, NI) and subjects’ chronological age (in month) were calculated in ROIs when a significant effect (i.e. above chance global decoding of action category) was found.

#### 2.8.3. Searchlight MVPA

To complete the results obtained with the ROI-based decoding and test the presence of additional putative brain areas for decoding Social vs Non-Social and Transitive vs Intransitive, we carried out a whole-brain searchlight analysis with 12mm radius spheres (about 463 voxels). MVPA classification was carried out with the same parameters and procedure as the ROI-based MVPA, within each sphere as the searchlight moved across the brain, and the classification accuracy was stored at the central voxel, yielding a 3D brain map of classification accuracy (Haynes, 2015). To identify regions where classification accuracy was significantly above chance (i.e., 50%) in adults and adolescents, the chance level was subtracted from classification maps, then these maps were normalized (MNI template) and smoothed (FWHM = 6mm). Then, we carried out one-sample t-tests for each group and each condition separately, corrected for multiple comparisons at the cluster level (FWE, *p* < .05).

## 3. Results

### 3.1. Behavioral ratings

We carried out one three-way ANOVA (Group x Sociality x Transitivity) separately for each rating (i.e., sociality and transitivity). Concerning the rating of the transitive dimension, we found a main effect of Transitivity *F*(1,49) = 176.88, *p* < .001, *η*_*p*_^*2*^ = .97, transitive actions (*M* = .93, *SD* = .32) were rated more transitive than intransitive videos (*M* = −.93, *SD* = .24), unsurprisingly. No other main effect nor interaction including the factor group were found. Concerning the rating of the social dimension, we found a main effect of Sociality *F*(1,49) = 623.82, *p* < .001, *η*_*p*_^*2*^ = .93, social videos (*M* = .79, *SD* = .51) were rated more social than the non-social actions (*M* = −.79, *SD* = .49). We also found a main effect of Transitivity *F*(1,49) = 11.32, *p* < .01, *η*_*p*_^*2*^ = .19, intransitive actions (*M* = .14, *SD* = 1.10) were rated more social than transitive actions (*M* = −.14, *SD* = .71). Finally, the ANOVA revealed an interaction between Sociality and Transitivity *F*(1,49) = 42.50, *p* < .001, *η*_*p*_^*2*^ = .46: there was no difference between non-social transitive (*M* = −.72, SD = .38) and non-social intransitive actions (*M* = − .86, SD = .58, *p* = .47), whereas social intransitive actions (*M* = 1.14, SD = .29) were rated more social than social transitive videos (*M* = .44, SD = .44, *p* < .001). No other main effect nor interaction including the factor group were significant.

### 3.2. Univariate fMRI results

We entered the individual maps of parameters estimates for the four action conditions (NI, NT, SI, ST) in a repeated-measure ANOVA with Sociality and Transitivity as within-subject factors and Age group as between-subject factor. The results are displayed in **Table 1** and **Figure 3**.

**Table 1.** Brain regions activated in the whole-brain analysis for the main effect of Age Group, Sociality and Transitivity

| Region Label |  | Extent | t-value | Peak MNI Coordinates |  |  |
| --- | --- | --- | --- | --- | --- | --- |
|  |  |  |  | *x* | *y* | *z* |
| **<u>Main effect of Group</u>** |  |  |  |  |  |  |
| *Adolescents > adults* |  |  |  |  |  |  |
| Ventral Medial Prefrontal Cortex | L | 58 | 5.94 | -6 | 61 | -8 |
| Temporo Parietal Junction | L | 116 | 4.49 | -43 | -59 | 25 |
| **<u>Main effect of Sociality</u>** |  |  |  |  |  |  |
| *Social/Non-Social* |  |  |  |  |  |  |
| Visual cortex | L/R | 8217 |  |  |  |  |
| Intracalcarine Cortex |  |  | 11.29 | 7 | -79 | 3 |
| Intracalcarine Cortex |  |  | 10.93 | -8 | -97 | 13 |
| Lingual Gyrus |  |  | 9.52 | -3 | -79 | -10 |
| Temporo-parietal Cortex | L | 1047 |  |  |  |  |
| pSTS/Middle Temporal Gyrus |  |  | 8.47 | -53 | -47 | 8 |
| Supramarginal Gyrus |  |  | 7.48 | -51 | -42 | 25 |
| Angular Gyrus |  |  | 6.95 | -56 | -62 | 10 |
| Temporo -parietal | R | 749 |  |  |  |  |
| pSTS/MTG |  |  | 7.97 | 47 | -42 | 10 |
| Supramarginal Gyrus |  |  | 6.33 | 67 | -39 | 23 |
| STS middle |  |  | 6.03 | 50 | -32 | -3 |
| Precuneus | L/R | 373 | 6.19 | -1 | -52 | 58 |
| Precentral Gyrus | L | 508 | 6.15 | -41 | -7 | 53 |
| Superior Frontal Gyrus |  |  | 5.02 | -26 | 4 | 60 |
| Pre-SMA |  |  | 4.49 | 12 | -4 | 63 |
| Precentral Gyrus | R | 114 | 5.69 | 47 | 1 | 55 |
| Superior Parietal Lobule | L | 78 | 4.97 | -33 | -49 | 35 |
| Inferior Frontal Gyrus | L | 55 | 4.37 | -46 | 14 | 23 |
| ***Non-Social /Social*** |  |  |  |  |  |  |
| Visual cortex | L/R | 8217 |  |  |  |  |
| Occipital Pole / Lateral Occipital |  |  | 12.61 | 32 | -92 | 5 |
| Occipital Pole / Lateral Occipital |  |  | 12.45 | -28 | -89 | 0 |
| Occipital fusiform gyrus |  |  | 12.35 | 17 | -87 | -8 |
| Temporal Occipital Fusiform Cortex |  |  | 11.29 | 27 | -49 | -18 |
| Anterior parietal cortex | L | 900 |  |  |  |  |
| Postcentral Gyrus/AIPS |  |  | 8.97 | -51 | -22 | 33 |
| Central Opercular Cortex |  |  | 6.62 | -56 | -17 | 18 |
| Superior Parietal Lobule |  |  | 6.45 | -28 | -47 | 68 |
| inferior Lateral Occipital Cortex | L | 92 | 7.55 | -46 | -69 | -8 |
| Precentral Gyrus | R | 76 | 5.04 | 30 | -12 | 58 |
| **<u>Main effect of Transitivity</u>** |  |  |  |  |  |  |
| ***Transitive / Intransitive*** |  |  |  |  |  |  |
| Medial occipital cortex | L/R | 10781 |  |  |  |  |
| Lingual gyrus |  |  | 16.50 | 15 | -87 | -10 |
| Lingual gyrus |  |  | 15.30 | -8 | -89 | -10 |
| Temporal Occipital Fusiform |  |  | 13.37 | 30 | -52 | -13 |
| Temporal Occipital Fusiform |  |  | 12.53 | -27 | -55 | -16 |
| Precentral Cortex | R | 450 | 10.23 | 25 | -7 | 53 |
| Precentral cortex | L | 300 | 7.30 | -23 | 1 | 55 |
| Superior Frontal sulcus | R | 74 | 5.70 | 22 | 21 | 40 |
| Parieto-occipital Cortex | L | 929 |  |  |  |  |
| Lateral occipital |  |  | 9.97 | -33 | -82 | 20 |
| Superior Parietal |  |  | 7.45 | -28 | -52 | 65 |
| Inferior temporal Cortex | R | 53 | 5.56 | 52 | -52 | -10 |
| Cerebellum (lobule VIII/ IX) | R | 83 | 5.45 | 15 | -47 | -50 |
| Cerebellum (lobule VIII/ IX) | L | 179 | 8.35 | -13 | -49 | -50 |
| Angular Gyrus | L | 57 | 4.68 | -48 | -62 | 23 |
| Posterior Cingulate Gyrus | R | 92 | 4.53 | 12 | -29 | 43 |
| *Intransitive > Transitive* |  |  |  |  |  |  |
| Medial occipital (early visual) cortex | L/R | 10781 |  |  |  |  |
| Cuneus |  |  | 14.16 | 12 | -94 | 18 |
| Cuneus |  |  | 12.65 | -11 | -99 | 8 |
| Intracalcarine Cortex |  |  | 9.92 | -3 | -77 | 10 |
| Lateral Occipital temporal cortex | R | 766 |  |  |  |  |
| Inferior Lateral Occipital Cortex (EBA) |  |  | 11.29 | 45 | -79 | -8 |
| Temporal Occipital Fusiform Cortex (FBA) |  |  | 7.50 | 45 | -44 | -20 |
| Posterior Superior Temporal Cortex | R | 527 |  |  |  |  |
| Supramarginal Gyrus |  |  | 7.09 | 52 | -37 | 8 |
| Post Superior Temporal Gyrus |  |  | 5.12 | 52 | -19 | -5 |
| Temporal pole | R | 77 | 6.61 | 37 | -4 | -45 |
| Temporal pole | L | 65 | 5.46 | -38 | -4 | -45 |
| Pericentral cortex | R | 313 |  |  |  |  |
| Central sulcus (hand area) |  |  | 5.28 | 35 | -19 | 40 |
| Central sulcus (index finger area) |  |  | 4.68 | 40 | -24 | 60 |
| Postcentral Cortex |  |  | 4.59 | 55 | -14 | 50 |

All results are thresholded at $p < .05$ (FWE corrected for multiple comparisons at the cluster level)

**Figure 3.**
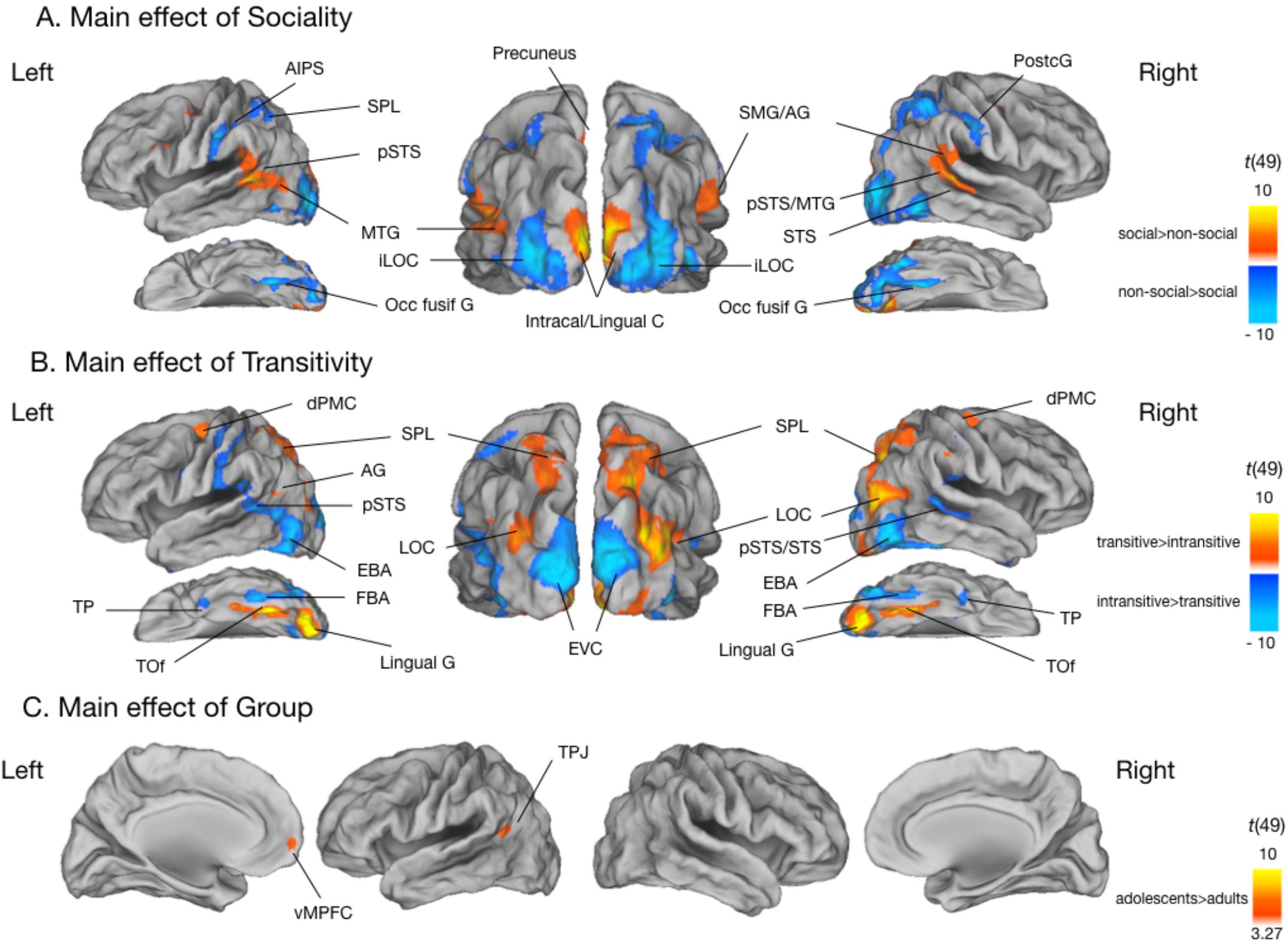
Brain activation associated with main effect of (A) Sociality; (B) Transitivity; and (3) Group. Activations are projected on PALS-B12 atlas surface configurations (Van Essen, 2005) : lateral fiducial surfaces. Statistical maps are FWE-corrected for multiple comparisons across the whole-brain at the cluster level; FWE, *p* < .05). AIPS: anterior intraparietal sulcus; SPL: superior parietal lobe; pSTS: posterior superior temporal sulcus; MTG: middle temporal gyrus; STS: superior temporal sulcus; iLOC: inferior lateral occipital cortex; Occ fusif G: occipital fusiform gyrus; Intracal: intracalcarine cortex; SMG: supramarginal gyrus; AG: angular gyrus; PostG: postcentral gyrus; dPMC: dorsal premotor cortex; LOC: lateral occipital cortex; TP: temporal pole; TOf: temporo-occipital fusiform gyrus; Lingual G: Lingual gyrus; EBA: extrastriate body area; FBA: fusiform body area; EVC: extrastriate visual cortex; vMPFC: ventral medial prefrontal cortex; TPJ: temporoparietal junction.

The ANOVA revealed a main effect of Sociality (see **Figure 3A**): observing social compared to non-social actions induced stronger activity in AON regions in bilateral posterior superior temporal sulcus and bilateral middle temporal gyrus, bilateral supramarginal gyrus, bilateral precentral gyrus, in left superior parietal lobe and in left inferior frontal gyrus bilateral, as well as in superior frontal gyrus, SMA, precuneus bilateral visual cortices (intracalcarine cortex and lingual gyrus). The reverse contrast yielded significant activation in left anterior parietal cortex (AIPS/SPL), left inferior occipital cortex and right precentral gyrus, as well as in occipital pole and lateral occipital cortex.

We found a main effect of Transitivity (see **Figure 3B**): observing transitive actions was associated with stronger activity in bilateral medial occipital cortex, bilateral precentral cortex, right superior frontal sulcus, left parieto-occipital cortex, right inferior temporal cortex, bilateral cerebellum (lobule VIII/IX), left angular gyrus and right posterior cingulate cortex. The reverse contrast revealed significant activations in bilateral early visual cortices (cuneus), right lateral occipital temporal cortex (EBA/FBA), right posterior superior temporal cortex (SMG/pSTS), bilateral temporal poles, right pericentral cortex (central sulcus and postcentral cortex).

There was also a main effect of Age group. The contrast adolescents versus adults revealed higher activation in adolescents, when observing action compared to the control condition activation, in left ventral medial prefrontal cortex and in left temporoparietal junction (**Figure 3C**).

We did not observe any significant interaction between Sociality and Transitivity in any region. Finally, the ANOVA did not reveal any interaction between the factors Sociality or Transitivity and Age group nor three-way interaction.

### 3.3 ROI MVPA

#### 3.3.1. Decoding social vs non-social and transitive vs intransitive actions

Significant above-chance decoding was found in all the regions of the AON, for both adolescents and adults. LOTC and IPS/SPL showed the highest decoding accuracies for the social dimension and LOTC for the transitive dimension (**Figure 4**). We also assessed the significance of decoding in these regions at the individual level using permutations (Etzel & Braver, 2013; see **Extended Data Figure 4-1**), with a cutoff of *p* < .05. For the social dimension, all adults (left = 100% and right = 100%) and nearly all adolescents (left = 93% and right = 96%) decoded significantly in the LOTC, but in IPS/SPL the proportion of adolescents (left = 70% and right = 44%), for whom decoding was significant, was lower than that of adults (left = 79% and right = 79%). For the transitive dimension, decoding was significant in all participants in the LOTC.

**Figure 4.**
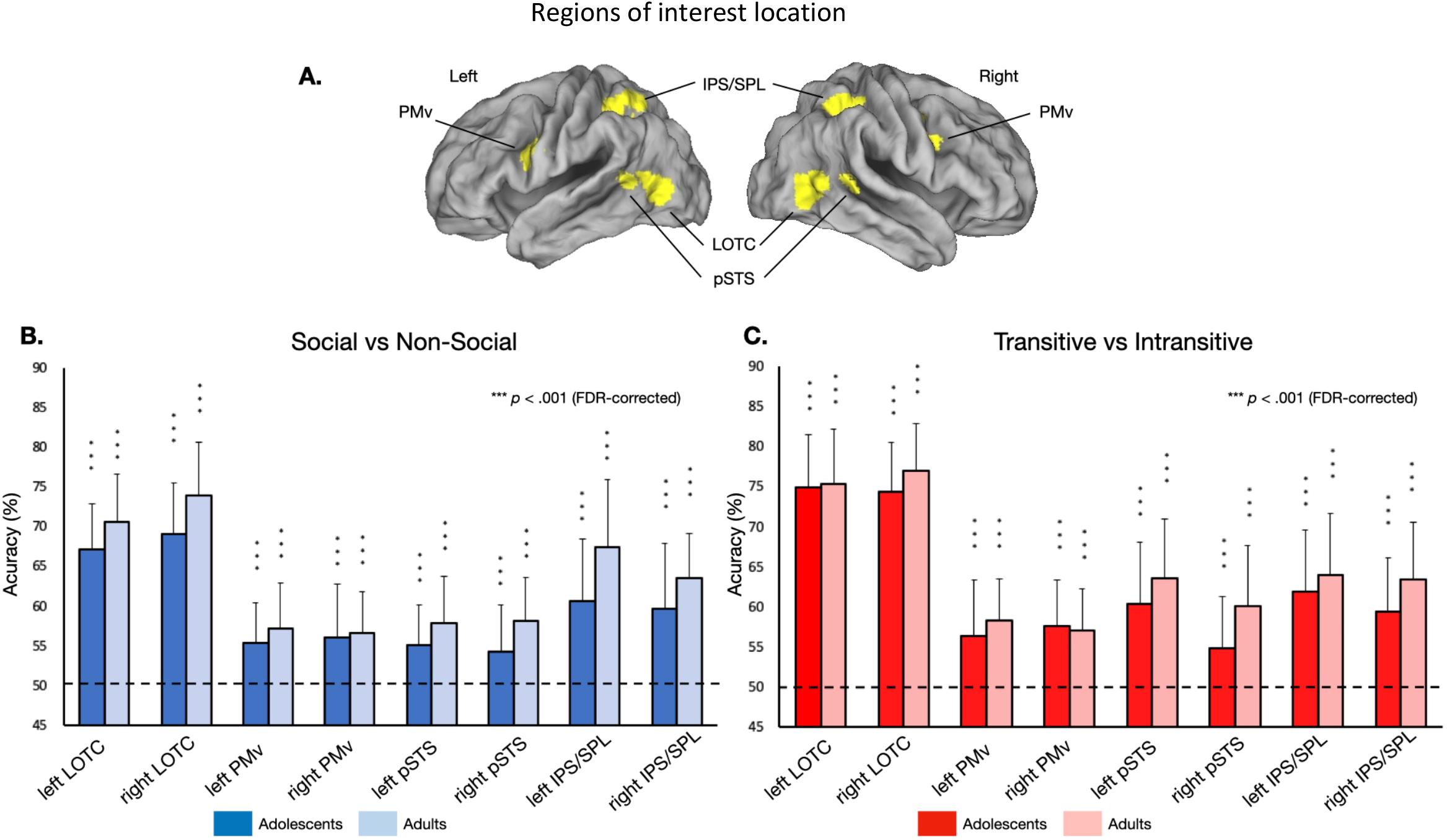
ROI MVPA results. (A) Illustration of the 8 functionally defined ROIs used in the present study derived from the meta-analysis of Grosbras et al. (2012). MNI-coordinates of the different ROIs are represented in **Extended Data Table 4-1**. (B) Group averaged decoding accuracies for (B) decoding social versus non-social (blue) and (C) transitive versus intransitive (red) actions for adolescents (dark) and adults (light). Error bars indicate Standard Deviation (SD). Asterisk represents statistical significance (FDR-corrected for the number of tests). Dotted line indicates decoding accuracy at chance-level (50%). Individual data is represented in **Extended data Figure 4-2**.

In a second step, we compared classification performance for adolescents and adults in LOTC, PMv, pSTS, and IPS/SPL, by entering mean classification accuracies in two-way ANOVAs with Hemisphere (Left, Right) as within subject factor and Age group (Adolescents, Adults) as between factor. These analyzes were performed for each dimension (i.e., transitivity and sociality) separately. Concerning the social dimension, the ANOVAs revealed a main effect of Age group in IPS/SPL, *F*(1,49) = 9.2, *p* < .01, in pSTS, *F*(1,49) = 8.17, *p* < .01, and in LOTC, *F*(1,49) = 7.23, *p* < .01, with higher decoding values for adults. There was a main effect of Hemisphere in LOTC, *F*(1,49) = 9.11, *p* < .01, and in IPS/SPL, *F*(1,49) = 4.07, *p* = .049, with higher decoding values in the right hemisphere. There was no interaction between Hemisphere and Age group (All *p* > .10). Concerning the transitive dimension, the ANOVAs revealed a main effect of Age group in pSTS, *F*(1,49) = 6.35, *p* = .015. There was a main effect of Hemisphere in pSTS, *F*(1,49) = 16.64, *p* < .001. There was no interaction between Hemisphere and Age group (All *p* > .10).

#### 3.3.2 Searchlight MVPA

Significant decoding was found for social and transitive actions bilaterally in brain areas typically associated with the AON including LOTC, PMv, pSTS, and IPS/SPL in both groups of participants (see **Figure 5A**). Moreover, when comparing accuracy maps for adults and adolescents using two-sample t-tests, we found significant clusters in bilateral IPS for Social versus Non-Social actions and in right pSTS for Transitive vs Intransitive actions (see **Figure 5B** and **Extended Data Table 5-1**), thus confirming the results obtained in the ROI analysis.

**Figure 5.**
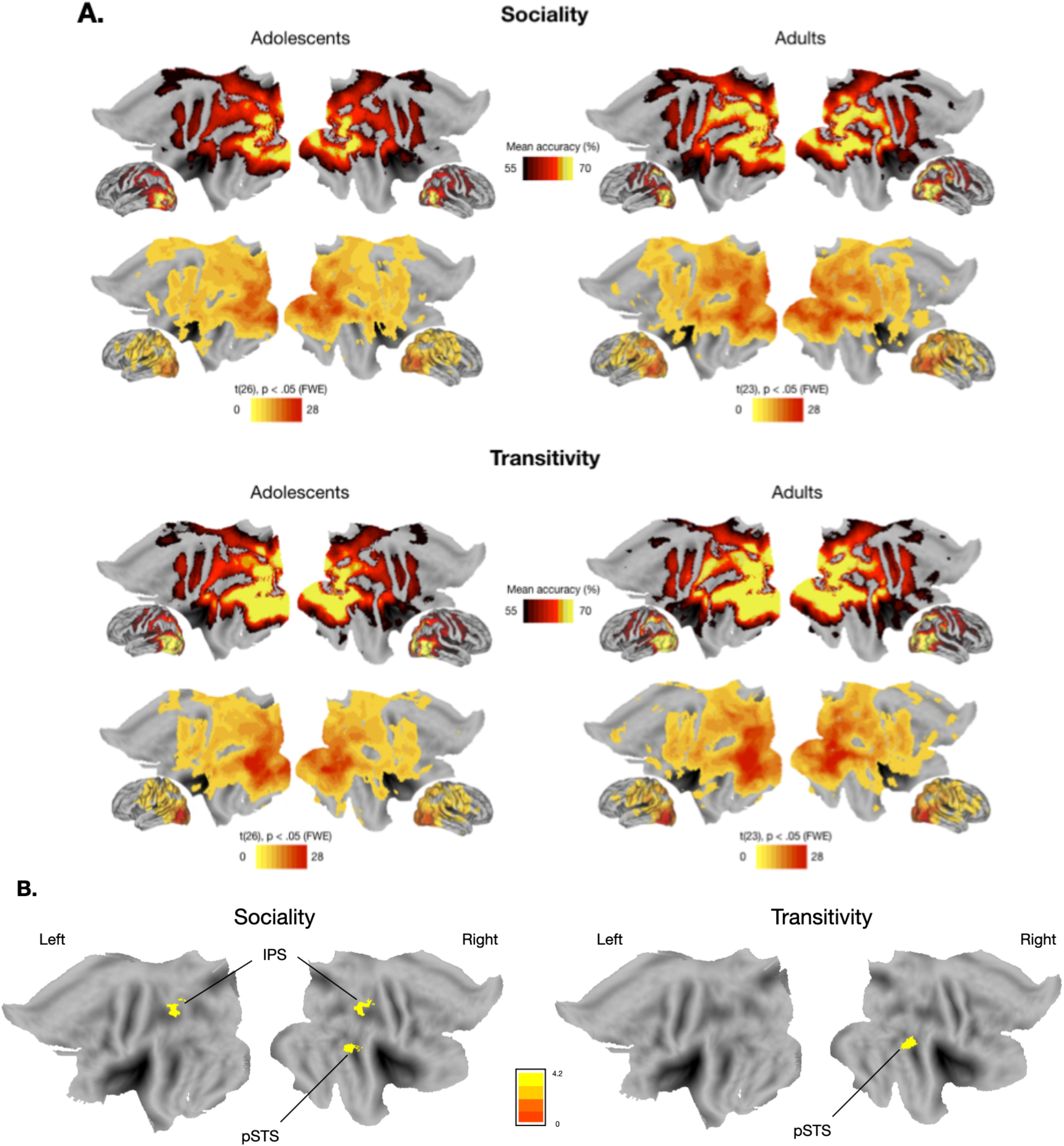
MVPA searchlight analyses. **(A)** Mean accuracy maps and statistical comparison maps of the searchlight decoding for Social versus Non-Social actions (chance level = 50%) and for Transitive versus Intransitive actions (chance level = 50%) for Adults and Adolescents. **(B)** Comparison of Searchlight accuracy maps of Adults and Adolescents using two-sample t-tests for Sociality and Transitivity separately. Corrections for multiple comparisons were applied at the cluster level (FWE, *p* < .05). Coordinates of significant clusters are presented in **Extended data Table 5-1**.

#### 3.3.3. Decoding individual action classes (NI, NT, SI, and ST)

We also carried out a four-way classification in each ROI and each participant and derived confusion matrices representing the pairwise decoding accuracies across conditions (i.e. how often a pattern corresponding to a condition is correctly decoded: matrix diagonal) and confounded with each of the other conditions (see **Figure 6**). The classifier was able to correctly discriminate each action class above chance in LOTC, IPS/SPL, and PMv and in a lesser extent in pSTS (see **Extended Data Figure 6-1**). The confusion matrices were highly similar between adults and adolescents.

**Figure 6.**
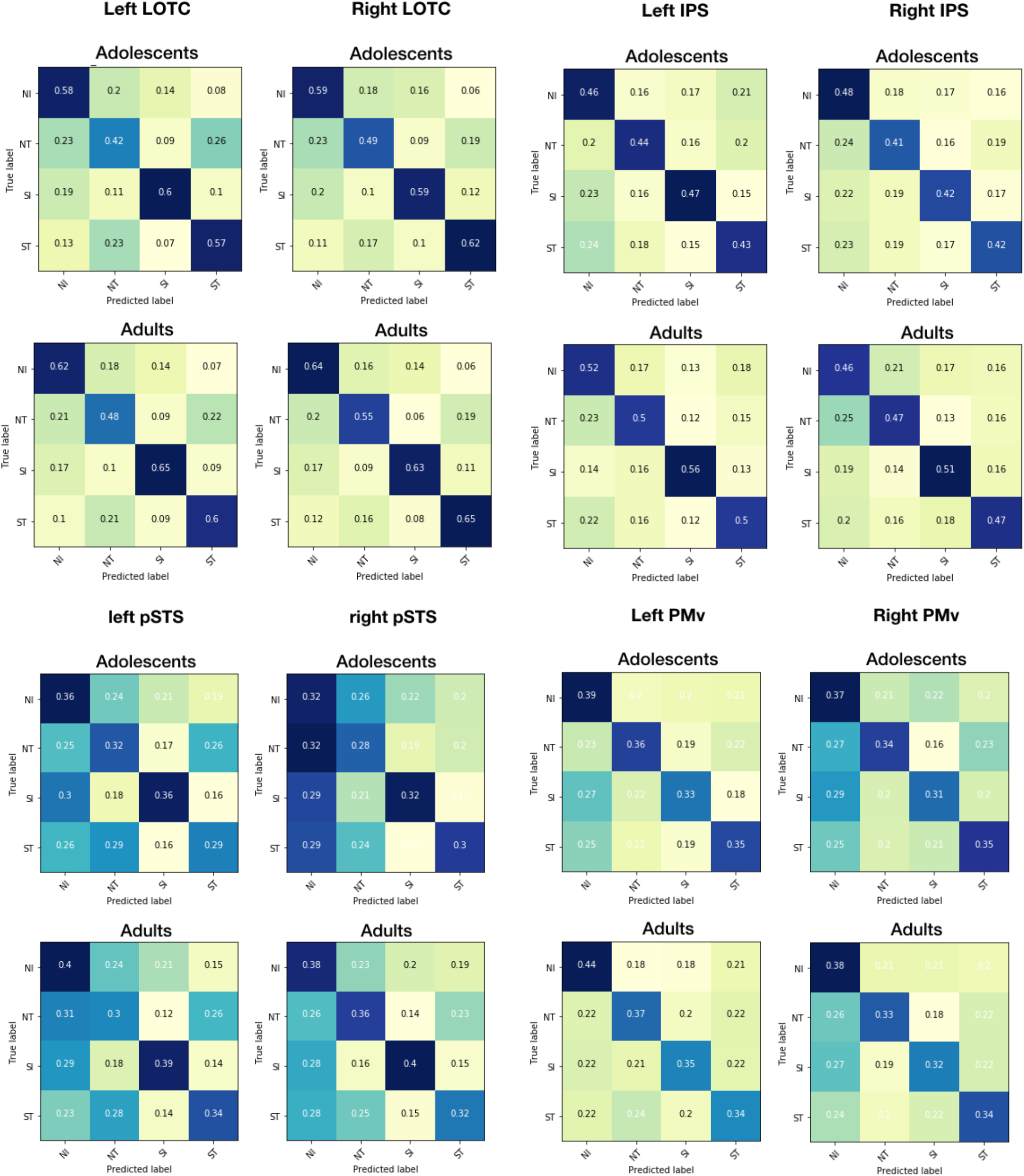
Confusion matrices for each action class in ROIs for adolescents and adults, providing the percentage of correct classifications (diagonals) and misclassifications (off diagonals). The lower the percentage, the more the cell is light-yellow colored and the higher the percentage, the more the cell is dark blue colored. Comparison of diagonal values to chance (0.5) are presented on **Extended data Figure 6-1**.

To investigate potential differences between adolescents and adults for each action class, mean classification accuracies were entered in ANOVAs with Sociality and Transitivity as within-subject factor and Age group as between-subject factor. Mean classification accuracies were averaged from the two hemispheres, as no interaction with the factor Hemisphere was significant in the first ROI MVPA. We carried out ANOVAs separately for each ROI (LOTC, PMV, IPS/SPL, and pSTS). These analyzes revealed a trend to significance for the interaction Age Group x Sociality x Transitivity in the IPS/SPL, *F*(1,49) = 3.90, *p=* 0.054 (**Figure 7A**), but this double interaction was not significant either in the LOTC, *F*(1,49) < 1, in PMv, *F*(1,49) < 1, or in the pSTS, *F*(1,49) < 1. In the IPS/SPL, decoding accuracies were higher for adults compared to adolescents for NT, *t*(49) = −2.10, *p* = .02, SI, *t*(49) = −2.41, *p* < .01, and ST, *t*(49) = −1.71, *p* = .047, but not for NI, *t*(49) = −.51, *p* = .31. Finally, we found a significant correlation between decoding accuracies and chronological age in IPS/SPL only in adolescents for SI, *r*(25) = .47, p = .012, and ST, *r*(25) = .52, p < .01 (see **Figure 7B**).

**Figure 7.**
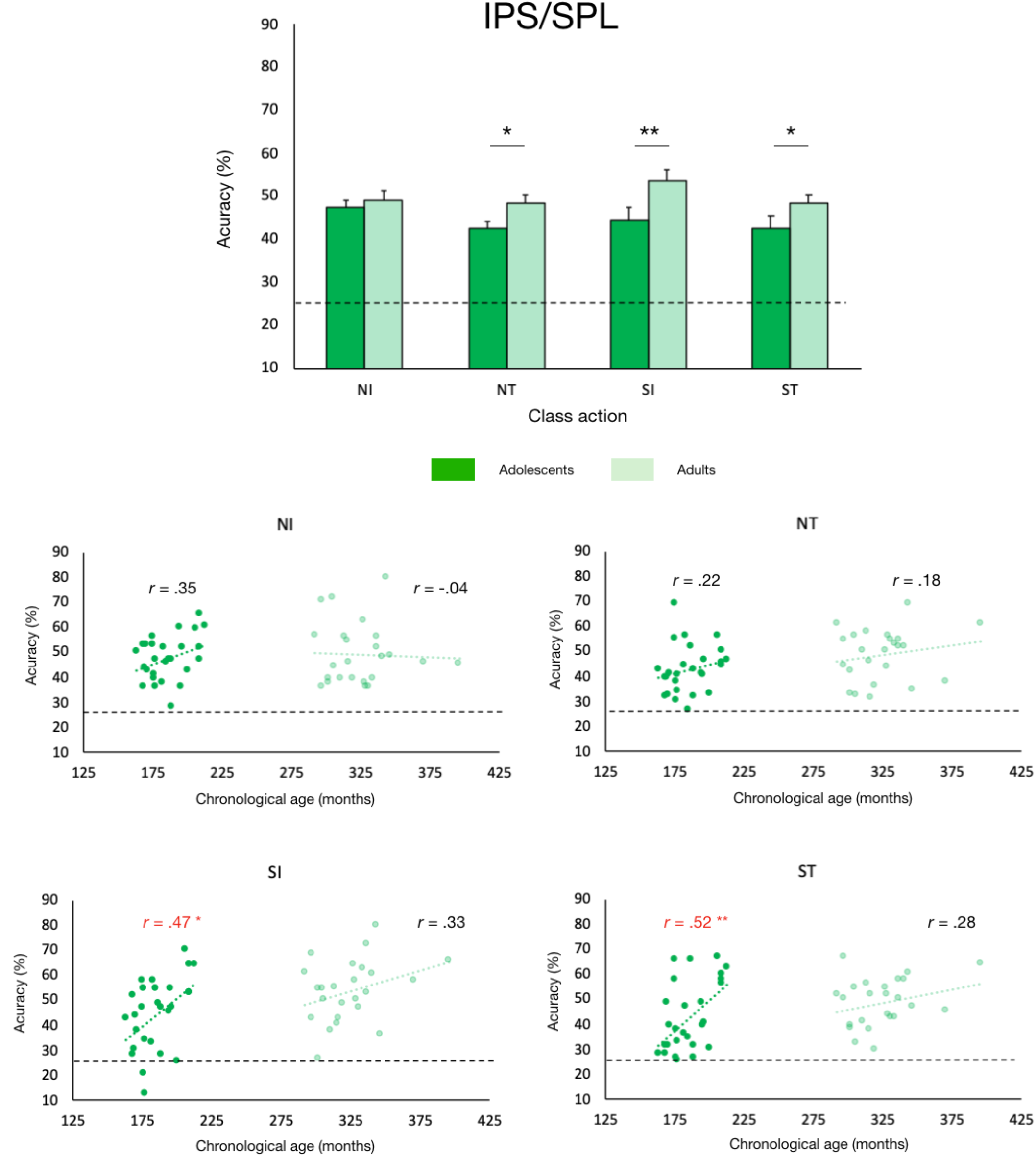
Mean decoding accuracies in IPS/SPL for each category of action for adolescents (dark) and adults (light). **Upper panel:** ANOVA on mean decoding accuracies with sociality and transitivity as within-subject factors and Age group as between-subject factor. **Bottom panels:** mean decoding accuracies are plotted against chronological age for each group and each class action (NI, NT, SI, and ST). Significant coefficient correlations (Pearson) are indicated in red. Dotted lines represent decoding accuracy at chance-level (25%). * *p* < .05, ** *p* < .01.

## 4. Discussion

Our univariate analyses indicate that all components of the AON are in place in adolescence and are engaged to the same level as in adults. Moreover, multivariate analyses showed that, like in adults, regions of this network contain information related to the content of actions. Yet this fine-grained action representation becomes more robust between adolescence and adulthood in IPS/SPL, pSTS and LOTC. Additionally, outside the AON we observed higher activity in adolescents in the MPFC and TPJ, two regions of the mentalizing network.

These findings extend previous reports of adult-like AON engagement in childhood and early adolescence (Ohnishi et al., 2004; Pokorny et al., 2015; Biagi et al., 2016; Morales et al., 2019) by testing advanced and post pubertal adolescents (14-17 years old). Furthermore, we show that the modulation of AON activity by the transitive and social dimensions of the observed actions is similar in adolescents and adults. Social actions induced higher activity than non-social actions in the pSTS, supramarginal gyrus, and precentral cortex, independently of whether these actions also involved an object. This complements previous adults studies that investigated either object-directed actions with a social intent or communicative symbolic actions or interactions (Iacoboni et al., 2004; Montgomery et al., 2007; Centelles et al., 2011; Saggar et al., 2014; Sliwa and Freiwald, 2017; Walbrin et al., 2018). In contrast, non-social actions engaged the most posterior parts of the temporal occipital cortex, as well as anterior parietal/post central areas, perhaps in relation to the fact that they drew attention to somato-sensation in the active actor, in particular in the stroking or rubbing videos. Observing transitive, relative to intransitive, actions yielded significant activation in bilateral medial fusiform gyrus, which is not typically included in the AON, but involved in processing information about objects (Mahon et al., 2007) and object-directed actions (Chen et al., 2016). We also observed bilateral activation of IPS/SPL and dPMC, which are part of a frontoparietal network involved in grasping and reaching (Daprati and Sirigu, 2006), as well as in observing others using tools (rev. in Reynaud, Navarro, Lesourd, & Osiurak, 2019). Observing intransitive versus transitive actions revealed activation in bilateral pSTS/STS and lateral occipitotemporal cortex (extending into the fusiform gyrus). This latter region is likely to encompass the extrastriate body area (EBA) and the fusiform body area (FBA), which selectively process visual features of human bodies (Downing & Peelen, 2011). Interestingly, Wagner and colleagues (2016), using naturalistic movie stimuli showed that FMRI signal peaks in the lateral fusiform gyrus occurred more frequently in response to scenes depicting a person (face or body) engaged in a social action, while peaks in the medial fusiform gyrus occurred for scenes with objects, landscapes or buildings, irrespective of the presence of social cues. In line with our data, this suggests that EBA and FBA are more engaged by intransitive than transitive actions stimuli and the reverse for the medial fusiform gyrus.

These findings are comforted by the multivariate analyses that provide evidence of representations of both the social and transitive dimensions of actions in all parts of the AON. Yet, while the univariate analysis did not show any difference between adolescents and adults, multivariate decoding accuracies were lower in adolescents in the LOTC, pSTS and IPS/SPL for social versus non-social actions and in pSTS for transitive versus intransitive actions.

The LOTC contains a mosaic of focal but overlapping regions selective for particular types of information (like hand posture, body shape, tools) that forms the components of action representations important for action understanding and social interpretation (for discussions see Lingnau & Downing, 2015; Wurm and Caramazza, 2019). Some authors have suggested that the LOTC forms the perceptual anchor of a pathway that extends into the superior temporal cortex and temporal parietal junction, a gradient along which increasingly rich representations of the posture, movements, actions, and mental states of other people are constructed (Carter and Huettel, 2013). Here we found higher decoding accuracy in adults only for social but not for transitive actions. This suggests that the role of this region for social action representation is still immature in adolescence.

We also found significant differences between adolescents and adults in representation of the social but not the transitive dimension in a region within the IPS/SPL. This region is part of the dorsal frontoparietal network involved in planning (Przybylski and Króliczak, 2017), action emulation (Ptak et al., 2017), observation and execution of manipulative actions (Dinstein, Hasson, Rubin, & Heeger, 2007; Ferri, Rizzolatti, & Orban, 2015; Lanzilotto et al., 2019; Orban, Ferri, & Platonov, 2019; Reynaud, Lesourd, Navarro, & Osiurak, 2016; Reynaud et al., 2019) and could also play a more general role in action understanding, and therefore in social interactions, by representing actor-object interactions at a higher level of abstraction (Tunik et al., 2007; Ramsey and Hamilton, 2010). Our results suggest that discriminating whether goal-directed actions have a social purpose is less efficient in IPS/SPL of adolescents and improves gradually, as indicated by the linear correlation between decoding accuracy and age in the adolescent group.

As adolescence is a period of major social development, from a behavioral and neural point of view (reviewed in Burnett et al. 2011), it is perhaps not surprising to observe differences in the representation of the social dimension of actions. The lower decoding performance for the transitive dimension in adolescents in the pSTS is however less expected, considering that the understanding of object manipulation is certainly well mastered at this age. Our data might thus indicate that action representation, at the perceptual level, subtending action categorization in the pSTS might still change in adolescence. It has to be noted however that, in the pSTS, the social/non-social actions discrimination accuracy was weaker compared to transitive/intransitive actions and also not as high as in LOTC or IPS/SPL, like in Wurm and colleagues (2017) study; at individual level the decoding was significant (permutation tests) on only about half of the adults and one-third of the adolescents. It is coherent with the interpretation that pSTS responds to mutual interactions between coacting agents (Isik et al., 2017; Walbrin et al., 2018): there was no mutual interactions between actors neither in Wurm and colleagues (2017) nor in our study (i.e., one acting agent and one passive agent). In any case the fact that we observed age differences for both the social and the transitive dimensions indicates that the representation of action categories in this region is still different from that of adults.

Age-differences emerged from our multivariate but not univariate analyses suggesting that different patterns of voxels may capture subtle changes between adolescents and adults that could not be revealed at the voxel-level. Differences in decoding accuracies between groups might be explained by different inter-subject variability (Bray et al., 2009). Individuals are maturing at different rates, and our adolescents’ sample is likely more heterogeneous than our adults’ sample. In our study, this explanation can apply for the right IPS/SPL where social versus non-social actions was decoded in only half of adolescents compared to 80% of adults (see Extended data Figure 4-2). Yet this is not the case in the other regions where higher decoding accuracy is observed in adults despite a similar proportion of adults and adolescents with significant decoding. This shows that interindividual variability in functional organization may account for only some but not all differences between adolescents and adults and that inter-subject variability decreases with age non-homogeneously in different AON parts.

Outside AON, the univariate analysis revealed that adolescents but not adults recruited the vMPFC and TPJ, two regions commonly attributed to the mentalizing network (Frith and Frith, 2007; Van Overwalle and Baetens, 2009), usually engaged when people make attributions about the mental states of others. Developmental studies reported that during such tasks, adolescents activated the MPFC to a greater extent than adults (reviewed in Blakemore, 2008). It may be that during our task, adolescents also inferred thoughts and intentions, independently of the transitive or social nature of the actions. Future studies should investigate behavioral correlates of viewing these actions as well as links between the AON and mentalizing areas.

In conclusion, our results contribute to the understanding of the AON development in adolescence. In line with our hypothesis, we revealed age differences in the local pattern of activation representing the social dimension of an action in LOTC, IPS/SPL and pSTS, as well as strengthening of the representation of the transitive dimension in the pSTS. We observed no evidence of differences in the precentral regions.This underlies adolescent development in the functional organization of the posterior parts of the AON. Future studies should investigate how other featural or contextual components of actions are represented in the AON of adolescents, in relation to changes in social perception skills.

## Data and Code availability statement

Unthresholded statistical maps for the main contrasts of interest can be visualized on NeuroVault (https://neurovault.org/collections/8403/). Behavioral and preprocessed neuroimaging data will be posted on a public repository (OpenfMRI) after publication of the research article. Stimulus materials and code are available upon reasonable request.

## Acknowledgements

This research was supported by grants from the Agence Nationale de la Recherche (France). ANR-14-ACHN-0023; ANR-16-CONV-0002 (ILCB) and the Excellence Initiative of Aix-Marseille University (A*MIDEX; ANR-11-IDEX-0001-02).

